# AMPK mediates early activation of the unfolded protein response through a positive feedback loop in palmitate-treated muscle cells

**DOI:** 10.1101/2021.10.12.464004

**Authors:** Jing Gong, Lu Wang, Wuchen Tao, Zonghan Liu, Xiangsheng Pang, Wenjiong Li, Yaxuan Liu, Xiaoping Chen, Peng Zhang

**Author notes:** **Correspondence:** Peng Zhang, Xiaoping Chen. These authors have contributed equally to this work and share first authorship (Jing Gong, Lu Wang).

## Abstract

Activation of the unfolded protein response (UPR) is closely associated with the pathogenesis of many metabolic diseases including obesity and type 2 diabetes. There is increasing evidence for the interdependence of the UPR and metabolic signaling pathways. The AMP-activated protein kinase (AMPK) signaling pathway controls energy balance in eukaryotes. The aim of this study was to investigate the possible interaction between AMPK signaling and the UPR in muscle cells exposed to a saturated fatty acid, as well as the underlying mechanism. The UPR was induced in C2C12 myotubes by treatment with palmitate along with activation of AMPK signaling. Inhibiting the AMPK pathway with compound C attenuated palmitate-induced UPR activation, while inhibiting the UPR with taurourdodeoxycholic acid alleviated palmitate-induced AMPK activation, suggesting a positive feedback loop between the UPR and AMPK. Additionally, 5-amino-1-β-D-ribofuranosylimidazole-4-carboxamide, an AMPK agonist, caused a dose- and time-dependent upregulation of genes related to the UPR, including activating transcription factor (ATF)4, binding immunoglobulin protein (BIP), and growth arrest and DNA damage-inducible protein (GADD)34. These results provide the first evidence for the involvement of AMPK signaling in the early activation of the UPR induced by saturated fatty acid in skeletal muscle, and suggest that physiologic or pharmacologic activation of the AMPK pathway (ie, by exercise or metformin, respectively) can promote skeletal muscle health and function and thus improve quality of life for individuals with metabolic disorder due to a high-fat diet or obesity.

## 1 Introduction

Adult skeletal muscle shows considerable plasticity that allows a rapid response under a variety of physiologic and pathologic conditions [1], which is facilitated by the sarcoplasmic reticulum, a specialized form of the endoplasmic reticulum (ER) [2]. Environmental or cell-intrinsic stimuli such as nutrient or oxygen deprivation, exposure to toxic substances, and oxidative stress can disrupt cellular homeostasis and induce ER stress, which leads to activation of the unfolded protein response (UPR) [3-5]. The canonical UPR in mammals is initiated by activation of 3 major ER transmembrane sensors—namely, PKR-like endoplasmic reticulum kinase (PERK), activating transcription factor (ATF)6, and inositol-requiring enzyme (IRE)1 [6-8]—that trigger the expression of downstream transcription factors (eg, ATF4, ATF6c, C/EBP homologous protein [CHOP], and spliced X-box binding protein 1 [XBP1s]). The main outcome of UPR signaling—specifically, of the early phase of the UPR—is the restoration of ER homeostasis through inhibition of protein synthesis or upregulation of ER chaperone proteins [9, 10]. However, prolonged UPR due to continuous stress can lead to the induction of apoptotic cell death [11, 12]. Thus, the UPR is a cellular mechanism that controls cell fate in response to stress.

ER stress and the UPR also can be activated in skeletal muscles exposed to metabolic stress, as occurs in diabetic patients [13] or by consumption of a high-fat diet [14, 15]. The high concentration of free fatty acids (especially saturated fatty acids [SFAs]) in plasma under these conditions is one of the main factors that trigger the UPR in skeletal muscle [16-18]. The UPR is closely associated with SFA-induced inflammation, autophagy, insulin resistance, and apoptosis in skeletal muscle [18-20], implying crosstalk between the UPR and signaling pathways that regulate metabolism [21, 22]. The AMP-activated protein kinase (AMPK) pathway, which is conserved across eukaryotes, integrates signals from multiple sources to control cellular energy balance. There is increasing evidence for the interaction of AMPK signaling with the UPR [23-30], but the mechanistic basis for the crosstalk between these two pathways has yet to be elucidated in different models of ER stress.

In this study we investigated whether there is crosstalk between AMPK signaling and the UPR induced by palmitate in skeletal muscle cells, as well as the possible underlying mechanism. We found that AMPK was activated in myotubes in response to treatment with the palmitate. Moreover, we showed that AMPK signaling crosstalks with early activation of the UPR via a positive feedback mechanism. Additionally, the AMPK agonist 5-amino-1-β-D-ribofuranosylimidazole-4-carboxamide (AICAR) induced mild UPR. These findings provide insight into the interactions between metabolic signals and homeostatic mechanisms in skeletal muscle cells that may be perturbed in metabolic disorders.

## 2 Materials and methods

### 2.1 Cell culture

Mouse C2C12 myoblast cells were cultured in high-glucose Dulbecco’s modified Eagle medium (DMEM) (Gibco, Grand Island, NY, USA) supplemented with 10% fetal bovine serum (FBS) (Sigma-Aldrich, St. Louis, MO, USA) and 1% antibiotics (100 U/ml penicillin and 0.1 mg/ml streptomycin; Gibco) at 37°C in a 5% CO2 atmosphere. When the cells reached 80%–90% confluence, the medium was replaced with high-glucose DMEM containing 2% horse serum (Gibco), which was changed daily. C2C12 myotubes were used for experiments after 5 days of differentiation.

### 2.2 Experimental treatments

Palmitate (Sigma-Aldrich; cat. no. P0500) was dissolved in ethanol and diluted to 500 μmol/l in DMEM containing 2% AlbumiNZ bovine serum albumin (MP Biomedicals, Solon, OH, USA; cat. no. 199896), 2% FBS (Atlanta Biologicals, Flowery Branch, GA, USA), 2 mmol/l L-carnitine (Sigma-Aldrich; cat. no. C0283), and 1% antibiotics [31]. Control cells were incubated in the same medium except that palmitate was substituted with an equal volume of ethanol. In some treatment conditions, 10 μmol/l compound C (prepared in dimethylsulfoxide [DMSO]; Sigma-Aldrich) was coincubated with palmitate for 12 h; DMSO was also used as a vehicle control for the treatments. To inhibit ER stress, C2C12 myotubes were pretreated for 1 h with 1 mM taurourdodeoxycholic acid (TUDCA) (Millipore, Billerica, MA, USA; cat. no. 580549) before adding palmitate for another 12 h. To activate AMPK signaling, AICAR (Sigma-Aldrich; cat. no. A9978) was added to the myotubes at a final concentration of 0.125–2 mmol/l for different times.

### 2.3 RNA extraction and real-time (RT-)PCR

Total RNA was extracted from C2C12 myotubes using TRIzol reagent (Invitrogen, Carlsbad, CA, USA; cat. no. 15596-026). RNA concentration and quality were verified using a Bio Photometer (Eppendorf, Hamburg, Germany). The PrimeScript RT reagent kit (Takara Bio, Otsu, Japan; cat. no. RR037A) was used to reverse transcribe total RNA (2 μg) into cDNA with random hexamer primers. RT-PCR was performed on a StepOnePlus RT-PCR system (Invitrogen) with fast SYBR Green Master Mix (Applied Biosystems, Foster City, CA, USA; cat. no. 4385612). Each RT-PCR mixture (final reaction volume=50 μl) contained 21 μl sterile water, 25 μl SYBR Green, 2 μl cDNA (500 ng/μl), and 1 μl each of forward and primers (10 pmol/μl). The reaction conditions were as follows: denaturation at 95°C for 10 s, annealing at the melting temperature of the specific primer set for 15 s, elongation at 72°C for 20 s, and a melting curve step. Target gene expression levels were normalized to that of the 18S rRNA gene. The following forward and reverse primers were used: BIP, 5′-AAACCAAGACATTTGCCCCAG-3′ and 5′-AGACACATCGAAGGTGCCG-3′; CHOP, 5′-CCTAGCTTGGCTGACAGAGG-3′ and 5′-CTGCTCCTTCTCCTTCATGC-3′; ATF4, 5′-GGAATGGCCGGCTATGG-3′ and 5′-TCCCGGAAAAGGCATCCT-3′; growth arrest and DNA damage-inducible protein (GADD)34, 5′-CGGAAGGTACACTTCGCTGA-3′ and 5′-CGGACTGTGGAAGAGATGGG-3′; XBP1s, 5′-GAGTCCGCAGCAGGTG-3′ and 5′-GTGTCAGAGTCCATGGGA-3′; unspliced XBP1 (XBP1u), 5′-AAGAACACGCTTGGGAATGG-3′ and 5′-ACTCCCCTTGGCCTCCAC-3′; and 18S rRNA, 5′-CCAGAGCGAAAGCATTTGCCAAGA-3′ and 5′-TCGGCATCGTTTATGGTCGGAACT-3′.

### 2.4 Immunoblotting

C2C12 myotubes were lysed in radioimmunoprecipitation assay buffer (Merck, Darmstadt, Germany) with complete EDTA-free protease and phosphatase inhibitors (Roche, Basel, Switzerland; cat. no. 04906845001). The supernatant was collected by centrifugation at 12,000×*g* for 10 min at 4°C and the protein concentration was determined using a microplate reader (Thermo Fisher Scientific, Waltham, MA, USA). Equal amounts of extracted protein (30 μg per lane) were denatured with gel loading buffer after centrifugation to remove insoluble material and separated by sodium dodecyl sulfate– polyacrylamide gel electrophoresis. The proteins were transferred to a nitrocellulose membrane that was blocked in 5% nonfat milk diluted in Tris-buffered saline with 0.1% Tween 20 (TBST) for 2 h and then incubated overnight at 4°C with primary antibodies against CHOP (cat. no. 5554), ATF4 (cat. no. 11815), AMPKα (cat. no. 2532), p-AMPKα (cat. no. 2531) (all from Cell Signaling Technology, Danvers, MA, USA), and β-actin (Santa Cruz Biotechnology, Santa Cruz, CA, USA; cat. no. sc130656). The following day, the membrane was washed 3 times with TBST and incubated for 2 h at room temperature with secondary antibodies in 5% nonfat milk, followed by incubation with enhanced chemiluminescence reagent (Thermo Fisher Scientific; cat. no. 34580) in a dark room. Protein bands were quantified using Image-Pro Plus v6.0 software (Media Cybernetics, Rockville, MD, USA); the intensity of the protein signal was normalized to that of β-actin.

### 2.5 Statistical analysis

Data are presented as mean ± SD. One-way analysis of variance followed by the Bonferroni posthoc test was used to compare the means of multiple groups using Prism v8.0 software (GraphPad, La Jolla, CA, USA). P≤0.05 was considered statistically significant.

## 3 Results

### 3.1 AMPK signaling is activated in the early stage of the UPR in myotubes

We investigated AMPK phosphorylation status and the expression of UPR markers in C2C12 myotubes treated with palmitate (a major component of dietary saturated fats) for different times. While total AMPKα levels remained constant over time, AMPKα phosphorylation was increased after 3 h of palmitate treatment, reaching a peak after 12 h; however, after 24 h, p-AMPKα level was lower than that in the control group (Figure 1A). The expression of UPR markers such as CHOP, ATF4, and XBP1s was also upregulated after 3 h of palmitate treatment and peaked at 12 h (Figure 1B). These results indicate that activation of AMPK signaling is closely associated with early activation of UPR induced by palmitate in myotubes.

**Figure 1.**
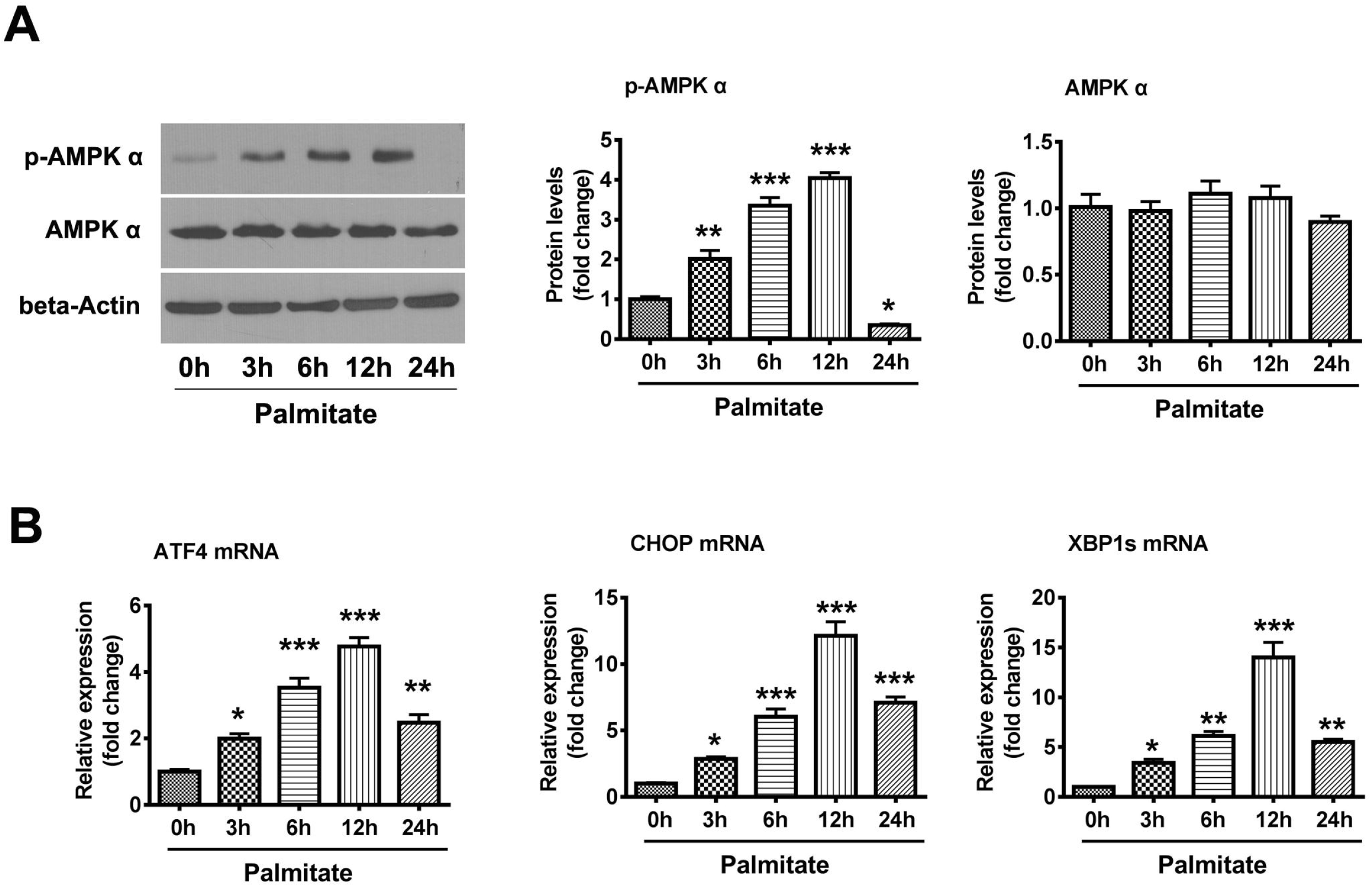
AMPK signaling is activated within 12 h of palmitate treatment. (A) C1C12 myotubes were incubated with 0.5 mM palmitate (Pal) for 0, 3, 6, 12, and 24 h. Proteins levels of AMPKα and p-AMPKα were evaluated by western blotting. The intensity of the protein bands was quantified by densitometry with Image-Pro Plus 6.0 software (n=4). (B) RT-PCR analysis of *ATF4, CHOP*, and *XBP1s* mRNA levels in C2C12 myotubes treated as described in panel A (n=4). Data are shown as the mean±SD. \**P*<0.05, \*\**P*<0.01, \*\*\**P*<0.001 vs control (0h) group (1-way analysis of variance).

### 3.2 AMPK signaling is involved in palmitate-induced UPR in myotubes

To clarify the interaction between the AMPK pathway and UPR, we treated C2C12 myotubes with palmitate for 12 h with or without compound C, a widely used specific inhibitor of AMPK. As expected, multiple factors involved in the UPR including ATF4, CHOP, GADD34, chaperone BIP, XBP1u, and XBP1s were upregulated by palmitate, as determined by RT-PCR (Fig. 2A); this was accompanied by increased AMPKα phosphorylation (Fig. 2B). AMPK inhibition by treatment with compound C abrogated the palmitate-induced upregulation of UPR markers (Fig. 2A). In agreement with the above findings, palmitate induced a marked increase in ATF4 and CHOP protein levels, which was partly abrogated by treatment with compound C (Fig. 2B). As a control, we confirmed that compound C completely abolished AMPK activation induced by palmitate (Fig. 2B). These data indicate that activation of AMPK signaling contributes to the early activation of the UPR induced by palmitate.

**Figure 2.**
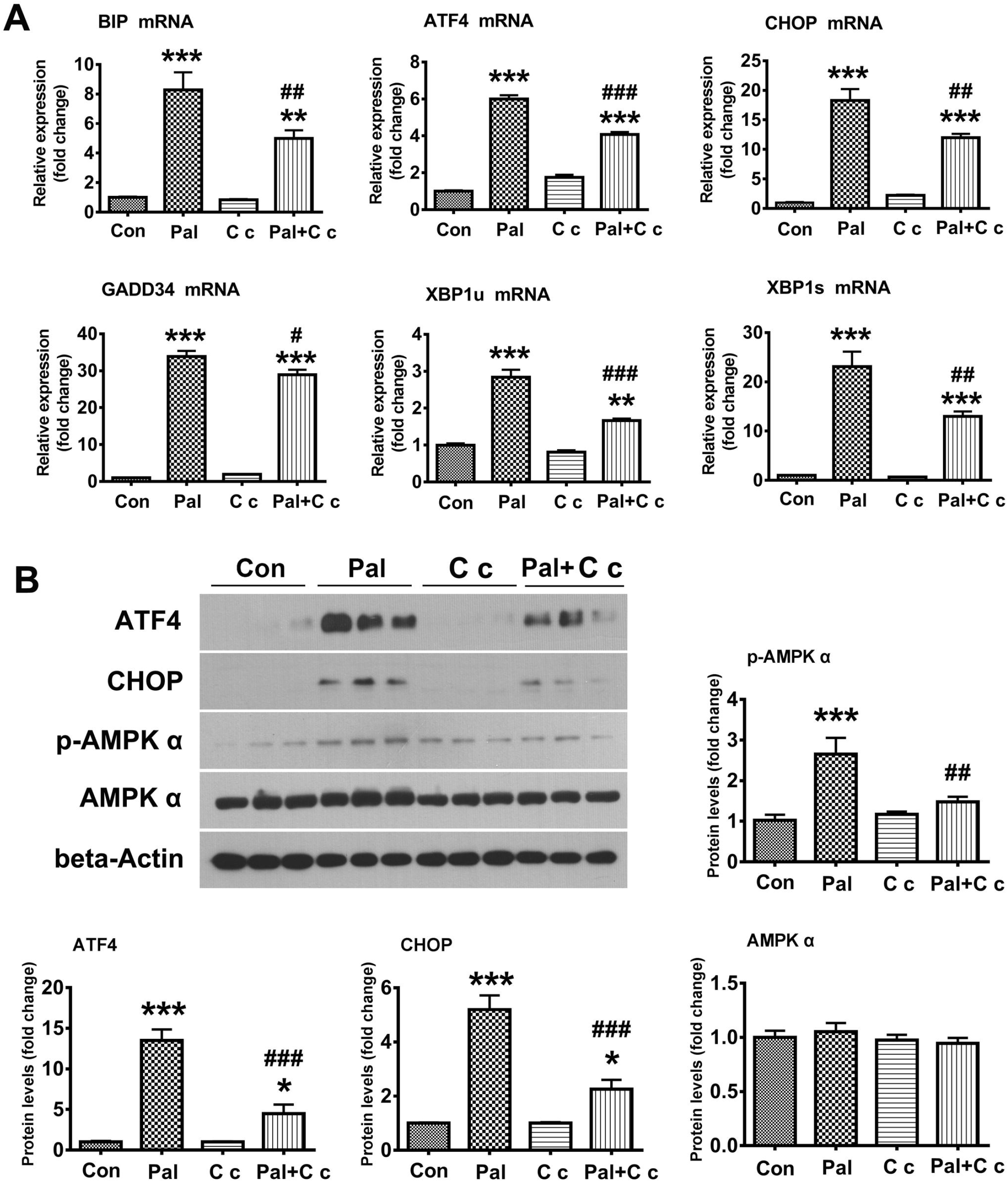
AMPK inhibition attenuates palmitate-induced UPR in C2C12 myotubes. (A) C2C12 myotubes were incubated for 12 h with 0.5 mM palmitate (Pal) in the presence or absence of 10 μM AMPK inhibitor compound C (C c). *BIP, ATF4, CHOP, GADD34, XBP1u*, and *XBP1s* mRNA levels were determined by RT-PCR (n=6). (B) Western blot analysis of AMPKα, p-AMPKα, CHOP, and ATF4 proteins levels in C2C12 myotubes treated as described in panel A. The intensity of the protein bands was quantified by densitometry with Image-Pro Plus 6.0 software (n=6). Data are shown as mean±SD. \**P*<0.05, \*\**P*<0.01, \*\*\**P*<0.001 vs control (Con) group; ^#^*P*<0.05, ^##^*P*<0.01, ^###^*P*<0.001 vs Pal group (1-way analysis of variance).

### 3.3 Inhibition of the UPR with TUDCA attenuates palmitate-induced AMPK activation

We further investigated whether inhibiting the UPR alters AMPK activation in C2C12 myotubes. The myotubes were pretreated with the UPR inhibitor TUDCA for 1 h before adding palmitate for 12 h. TUDCA significantly attenuated the palmitate-induced upregulation of ATF4 and CHOP (Fig. 3), confirming the pharmacologic inhibition of the UPR. Interestingly, TUDCA also abolished palmitate-induced AMPKα phosphorylation (Fig. 3), suggesting a positive feedback loop between the UPR and AMPK pathway in the early stage of palmitate treatment in muscle cells.

**Figure 3.**
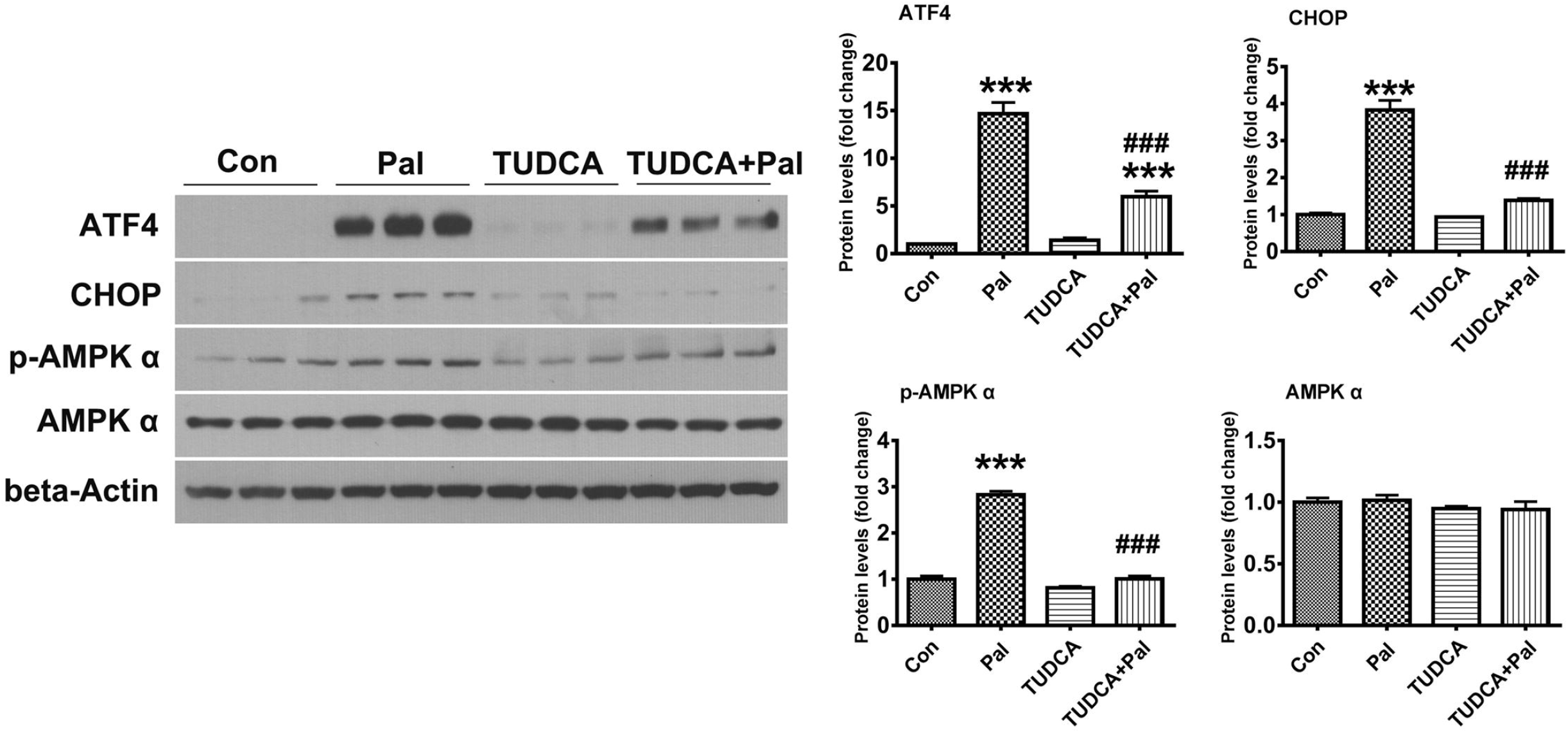
TUDCA alleviates palmitate-induced AMPK activation in C2C12 myotubes. C2C12 myotubes were pretreated for 1 h with 1 mM TUDCA or left untreated before palmitate (Pal) was added for another 12 h. Protein levels of AMPKα, p-AMPKα, ATF4, and CHOP were evaluated by western blotting (n=4). The intensity of the protein bands was quantified by densitometry with Image-Pro Plus 6.0 software (n=4). Data are shown as mean±SD. \*\*\**P*<0.001 vs control (Con) group; ^###^*P*<0.001 vs Pal group (1-way analysis of variance).

### 3.4 Pharmacologic activation of AMPK signaling induces mild UPR

Given our finding that AMPK activation contributes to palmitate-induced UPR, we speculated that pharmacologic activation of AMPK would be sufficient to induce the UPR in C2C12 myotubes. To test our hypothesis, C2C12 myotubes were treated with the AMPK agonist AICAR at concentrations ranging from 0.125–2 mM for 12 h to activate AMPK signaling. AMPK phosphorylation increased with AICAR concentration and was highest at 1 mM AICAR (Fig. 4A). Meanwhile, ATF4, GADD34, and BIP were upregulated by AICAR in a dose-dependent manner at concentrations <1 mM (Fig. 4B). A higher concentration of AICAR (2 mM) failed to induce AMPK activation to a greater extent than 1 mM, and the same was true for ATF4 and BIP expression (Fig. 4B). However, GADD34 was further upregulated whereas CHOP and XBP1u were downregulated by treatment with 2 mM AICAR (Fig. 4B). We also examined the effect of incubation time with AICAR (1 mM) on the UPR. After 6 h of treatment, there was significant activation of AMPK (Fig. 4C) while ATF4, GADD34, and BIP expression increased over time and was significantly higher after 12 h (Fig. 4D). These data indicate that pharmacologic activation of AMPK is sufficient to induce the UPR in myotubes—specifically ATF4, GADD34, and BIP—in a dose- and time-dependent manner.

**Figure 4.**
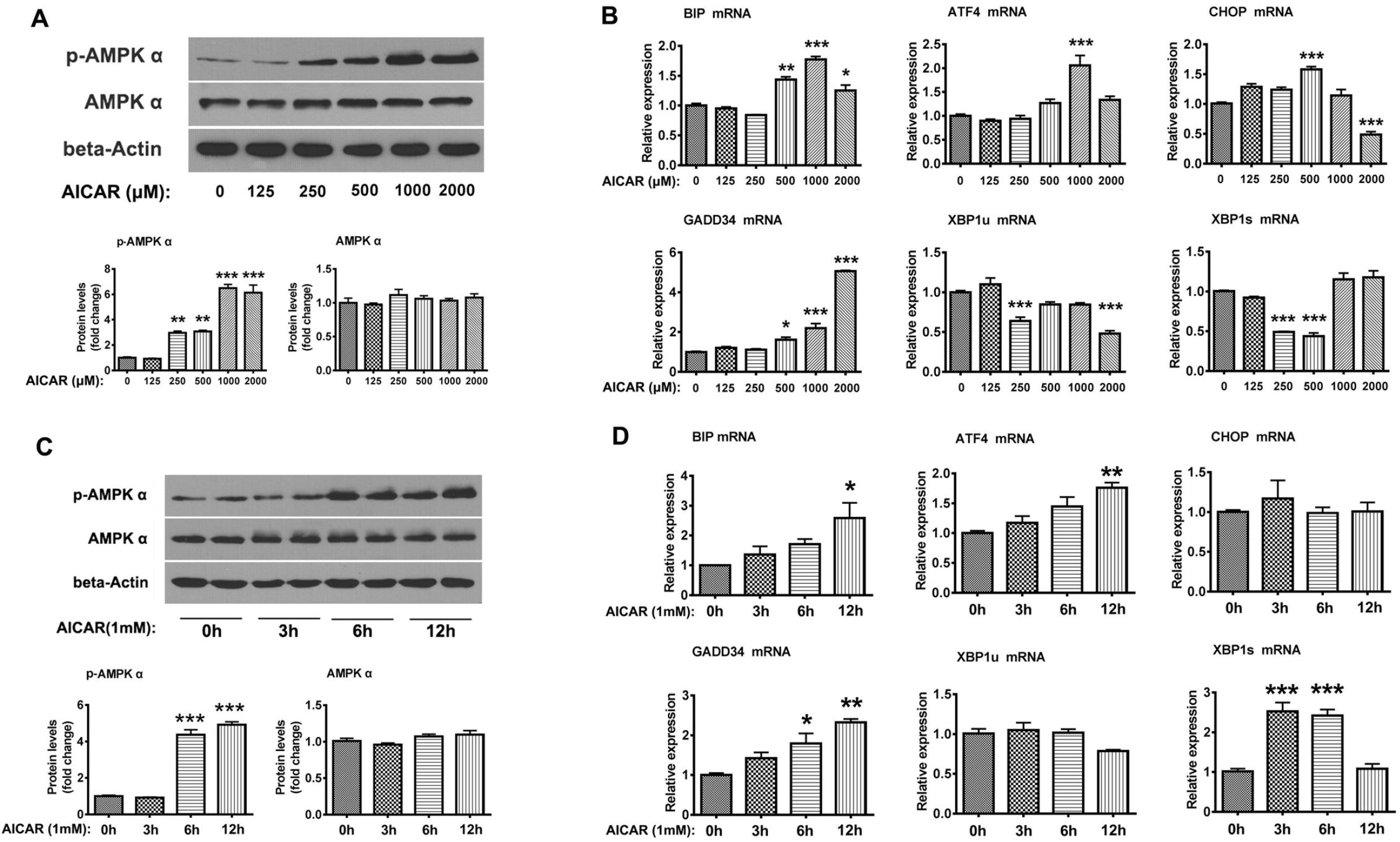
AMPK activation is sufficient to induce the UPR. (A) C2C12 myotubes were treated with different concentrations of AICAR (0.125–2 mM) for 12 h. Proteins levels of AMPKα, p-AMPKα, and ATF4 were determined by western blotting. The intensity of the protein bands was quantified by densitometry with Image-Pro Plus 6.0 software (n=4). (B) RT-PCR analysis of *BIP, ATF4, CHOP, GADD34, XBP1u*, and *XBP1s* mRNA levels in C2C12 myotubes treated as described in panel A (n=3). (C) C2C12 myotubes were treated with 1 mM AICAR for different times (0–12 h). Protein levels of AMPKα and p-AMPKα were determined by western blotting. The intensity of the protein bands was quantified by densitometry with Image-Pro Plus 6.0 software (n=4). (D) RT-PCR analysis of *BIP, ATF4, CHOP, GADD34, XBP1u*, and *XBP1s* mRNA levels in C2C12 myotubes treated as described in panel C (n=3). Data are shown as mean±SD. \**P*<0.05, \*\**P*<0.01, \*\*\**P*<0.001 vs control (0μM or 0h) group (1-way analysis of variance).

## 4 Discussion

The results of this study provide novel evidence for the interaction between the AMPK pathway and UPR in muscle cells exposed to palmitate, a major component of dietary saturated fats [32]. Specifically, we first observed unexpected activation of AMPK signaling within 12 h of palmitate treatment, which was accompanied by acute induction of the UPR. In support of our findings, a previous report showed that peroxisome proliferator-activated receptor gamma coactivator (PGC)-1α—one of the main downstream target genes of the AMPK pathway—was transiently upregulated after 4 and 8 h of palmitate treatment [33]. However, another study found that the AMPK pathway was inhibited in cells treated with palmitate for 16 h [18]. We speculate that this discrepancy is due to differences in treatment duration, especially given that changes induced by palmitate occur much more rapidly and are more dramatic in vitro than those observed in clinical obesity or induced by a high-fat diet. In fact, we also found that p-AMPKα was downregulated in cells exposed to palmitate for 24 h.

The association between the UPR and AMPK has been previously investigated [24, 34-37]. Some studies on palmitate-induced ER stress have demonstrated an inhibitory effect of AMPK signaling on the UPR in different tissues and cells [24, 34, 37]. For instance, pharmacologic activation of AMPK with AICAR was shown to suppress palmitate-induced ER stress in rat vascular endothelial cells [27](Li et al., 2015). In C2C12 myotubes, both GW501516 (a peroxisome proliferator-activated receptor [PPAR]δ receptor agonist) and oleate blocked palmitate-induced ER stress through an AMPK-dependent mechanism [18, 38]. Similarly, 5-lipoxygenase protected C2C12 myotubes from palmitate-induced ER stress via AMPK activation [39]. AMPK activation with AICAR (2 mM) effectively attenuated palmitate-induced ER stress in muscle cells [24]. However, our data showed that inhibiting the AMPK pathway with compound C attenuated the UPR induced by palmitate in myotubes, demonstrating a stimulatory effect of the AMPK pathway on palmitate-induced UPR. In agreement with our findings, the antidiabetic drug phenformin was shown to activate ER stress in an AMPK-dependent manner, while AMPK deficiency completely abolished phenformin-induced UPR [40]. Similarly, it was reported that AMPK activation induced mild UPR in C3H10T1/2 mouse mesenchymal stem cells [28]. We also demonstrated that inhibiting the UPR mitigated palmitate-induced AMPK activation, indicating a positive feedback loop between AMPK and the UPR in the early stage of palmitate treatment in muscle cells. This is the first report of a positive feedback regulation mechanism between the AMPK pathway and the UPR.

Interestingly, we also found that pharmacologic AMPK activation with AICAR was sufficient to induce the upregulation of UPR components in myotubes. In line with this finding, PGC-1α was shown to induce the expression of a variety of UPR-related genes in skeletal muscle [29]. Moreover, ER stress markers (eg, ATF3 and CHOP) and chaperones (eg, BIP and GRP94) were significantly unregulated in gastrocnemius muscle from transgenic mice with muscle-specific overexpression of PGC-1α [34]. PGC-1α overexpression also induced the expression of genes related to protein folding and the response to unfolded proteins in primary myotubes [6]. The increased expression of BIP and GADD34 caused by exercise was abolished in muscle-specific PGC-1α knockout mice, demonstrating that PGC-1α is important for the UPR in skeletal muscle [15]. Given the essential role of PGC-1α as an effector of the AMPK signaling pathway and its upregulation in the period soon after palmitate treatment [33], we speculate that PGC-1α is involved in the early activation of palmitate-induced ER stress in skeletal muscle.

In summary, we reported here the bidirectional crosstalk between AMPK signaling and early activation of the UPR in myotubes exposed to an SFA. We also showed that pharmacologic activation of AMPK was sufficient to induce a mild UPR in skeletal muscle cells. Our findings demonstrate an essential role for the AMPK pathway in restoring ER homeostasis through activation of the UPR in response to metabolic stress, which can guide the development of new strategies for the treatment of diseases such as obesity and diabetes through improvement of skeletal muscle metabolism.

## 6 Author Contributions

PZ and XC conceived the project and designed the study. JG, LW, ZL, XP, WT, and WL carried out the experiments. PZ, JG, WL, and YL analyzed the data. PZ, JG, LW, and XC wrote the manuscript. All authors read and approved the final version of the manuscript.

## 8 Acknowledgments

Not applicable.

## 10 Data Availability Statement

The datasets generated in this study can be obtained from the corresponding author on reasonable request.

## References

[1] Bohnert KR, McMillan JD, Kumar A. Emerging roles of ER stress and unfolded protein response pathways in skeletal muscle health and disease. J Cell Physiol. 2018;233:67–78. https://doi.org/10.1002/jcp.25852.

[2] Gallot YS, Bohnert KR. Confounding Roles of ER Stress and the Unfolded Protein Response in Skeletal Muscle Atrophy. Int J Mol Sci. 2021;22:2567. https://doi.org/10.3390/ijms22052567.

[3] Maekawa H, Inagi R. Stress Signal Network between Hypoxia and ER Stress in Chronic Kidney Disease. Front Physiol. 2017;8:74. https://doi.org/10.3389/fphys.2017.00074.

[4] Qi M, Dang Y, Xu Q, Yu L, Liu C, Yuan Y, et al. Microcystin-LR induced developmental toxicity and apoptosis in zebrafish (Danio rerio) larvae by activation of ER stress response. Chemosphere. 2016;157:166–73. https://doi.org/10.3389/10.1016/j.chemosphere.2016.05.038.

[5] Szpigel A, Hainault I, Carlier A, Venteclef N, Batto AF, Hajduch E, et al. Lipid environment induces ER stress, TXNIP expression and inflammation in immune cells of individuals with type 2 diabetes. Diabetologia. 2018;61:399–412. https://doi.org/10.1007/s00125-017-4462-5.

[6] Liu CY, Hsu CC, Huang TT, Lee CH, Chen JL, Yang SH, et al. ER stress-related ATF6 upregulates CIP2A and contributes to poor prognosis of colon cancer. Mol Oncol. 2018;12:1706–1717. https://doi.org/10.1002/1878-0261.12365.

[7] Ma Y, Brewer JW, Diehl JA, Hendershot LM. Two distinct stress signaling pathways converge upon the CHOP promoter during the mammalian unfolded protein response. Journal of Molecular Biology. 2002;318:1351–1365. https://doi.org/10.1016/s0022-2836(02)00234-6.

[8] Wang P, Li J, Tao J, Sha B. The luminal domain of the ER stress sensor protein PERK binds misfolded proteins and thereby triggers PERK oligomerization. J Biol Chem. 2018;293:4110–4121. https://doi.org/10.1074/jbc.RA117.001294.

[9] Rutkowski DT, Hegde RS. Regulation of basal cellular physiology by the homeostatic unfolded protein response. J Cell Biol. 2010;189:783–794. https://doi.org/10.1083/jcb.201003138.

[10] Wang BC, Zhang ST, Chen G. Research Progress of the UPR Mechanism and its Effect on Improving Foreign Protein Expression. Protein Pept Lett. 2020;27:831–40. https://doi.org/10.2174/0929866527666200407113549.

[11] Pinkaew D, Chattopadhyay A, King MD, Chunhacha P, Liu ZH, Stevenson HL, et al. Fortilin binds IRE1 alpha and prevents ER stress from signaling apoptotic cell death. Nature Communications. 2017;8:18. https://doi.org/10.1038/s41467-017-00029-1.

[12] Zhu Y, Xie M, Meng Z, Leung LK, Chan FL, Hu X, et al. Knockdown of TM9SF4 boosts ER stress to trigger cell death of chemoresistant breast cancer cells. Oncogene. 2019;38:5778–5791. https://doi.org/10.1038/s41388-019-0846-y.

[13] Liu J, Wu X, Franklin JL, Messina JL, Hill HS, Moellering DR, et al. Mammalian Tribbles homolog 3 impairs insulin action in skeletal muscle: role in glucose-induced insulin resistance. Am J Physiol Endocrinol Metab. 2010;298:E565–576. https://doi.org/10.1152/ajpendo.00467.2009.

[14] Sun JL, Park J, Lee T, Ji HJ, Jung TW. DEL-1 ameliorates high-fat diet-induced insulin resistance in mouse skeletal muscle through SIRT1/SERCA2-mediated ER stress suppression. Biochem Pharmacol. 2019;171:113730. https://doi.org/10.1016/j.bcp.2019.113730.

[15] Varone E, Pozzer D, Di Modica S, Chernorudskiy A, Nogara L, Baraldo M, et al. SELENON (SEPN1) protects skeletal muscle from saturated fatty acid-induced ER stress and insulin resistance. Redox Biol. 2019;24:101176. https://doi.org/10.1016/j.redox.2019.101176.

[16] Ebersbach-Silva P, Poletto AC, David-Silva A, Seraphim PM, Anhe GF, Passarelli M, et al. Palmitate-induced Slc2a4/GLUT4 downregulation in L6 muscle cells: evidence of inflammatory and endoplasmic reticulum stress involvement. Lipids Health Dis. 2018;17:64. https://doi.org/10.1186/s12944-018-0714-8.

[17] Peng G, Li L, Liu Y, Pu J, Zhang S, Yu J, et al. Oleate blocks palmitate-induced abnormal lipid distribution, endoplasmic reticulum expansion and stress, and insulin resistance in skeletal muscle. Endocrinology. 2011;152:2206–2218. https://doi.org/10.1210/en.2010-1369.

[18] Salvadó, L, Coll T, Gómez-Foix AM, Salmerón, E, Barroso E, Palomer X, et al. Oleate prevents saturated-fatty-acid-induced ER stress, inflammation and insulin resistance in skeletal muscle cells through an AMPK-dependent mechanism. Diabetologia. 2013;56:1372–1382. https://doi.org/10.1007/s00125-013-2867-3.

[19] Liong S, Lappas M. Endoplasmic reticulum stress regulates inflammation and insulin resistance in skeletal muscle from pregnant women. Mol Cell Endocrinol. 2016;425:11–25. https://doi.org/10.1016/j.mce.2016.02.016.

[20] Reddy SS, Shruthi K, Joy D, Reddy GB. 4-PBA prevents diabetic muscle atrophy in rats by modulating ER stress response and ubiquitin-proteasome system. Chem Biol Interact. 2019;306:70–77. https://doi.org/10.1016/j.cbi.2019.04.009.

[21] Yang BW, Qin Q, Xu L, Lv X, Liu Z, Song E, Song Y. Polychlorinated Biphenyl Quinone Promotes Atherosclerosis through Lipid Accumulation and Endoplasmic Reticulum Stress via CD36. Chem Res Toxicol. 2020;33:1497–1507. https://doi.org/10.1021/acs.chemrestox.0c00123.

[22] Yao S, Tian H, Miao C, Zhang DW, Zhao L, Li Y, et al. D4F alleviates macrophage-derived foam cell apoptosis by inhibiting CD36 expression and ER stress-CHOP pathway. Journal of Lipid Research. 2015;56:836–847. https://doi.org/10.1194/jlr.M055400.

[23] Behera S, Kapadia B, Kain V, Alamuru-Yellapragada NP, Murunikkara V, Kumar ST, Babu PP, et al. ERK1/2 activated PHLPP1 induces skeletal muscle ER stress through the inhibition of a novel substrate AMPK. Biochimica Et Biophysica Acta Molecular Basis of Disease Bba. 2018;186:1702–1716. https://doi.org/10.1016/j.bbadis.2018.02.019.

[24] Huang Y, Li Y, Liu Q, Zhang J, Zhang Z, Wu T, et al. Telmisartan attenuates obesity-induced insulin resistance via suppression of AMPK mediated ER stress. Biochem Biophys Res Commun. 2020;523:787–794. https://doi.org/10.1016/j.bbrc.2019.12.111.

[25] Hwang HJ, Jung TW, Choi JH, Lee HJ, Chung HS, Seo JA, et al. Knockdown of sestrin2 increases pro-inflammatory reactions and ER stress in the endothelium via an AMPK dependent mechanism. Biochim Biophys Acta Mol Basis Dis. 2017;1863:1436–44. https://doi.org/10.1016/j.bbadis.2017.02.018.

[26] Jung TW, Kim HC, Abd El-Aty AM, Jeong JH. Maresin 1 attenuates NAFLD by suppression of endoplasmic reticulum stress via AMPK-SERCA2b pathway. J Biol Chem. 2018;293:3981–3988. https://doi.org/10.1074/jbc.RA117.000885.

[27] Li J, Wang Y, Wang Y, Wen X, Ma XN, Chen W, et al. Pharmacological activation of AMPK prevents Drp1-mediated mitochondrial fission and alleviates endoplasmic reticulum stress-associated endothelial dysfunction. Journal of Molecular & Cellular Cardiology. 2015;86:62–74. https://doi.org/10.1016/j.yjmcc.2015.07.010.

[28] Son HE, Min HY, Kim EJ, Jang WG. Fat Mass and Obesity-Associated (FTO) Stimulates Osteogenic Differentiation of C3H10T1/2 Cells by Inducing Mild Endoplasmic Reticulum Stress via a Positive Feedback Loop with p-AMPK. Molecules and Cells. 2020;43:58–65. https://doi.org/10.14348/molcells.2019.0136.

[29] Wu J, Ruas JL, Estall JL, Rasbach KA, Choi JH, Li Y, et al. The unfolded protein response mediates adaptation to exercise in skeletal muscle through a PGC-1α/ATF6α complex. Cell Metabolism. 2011;13:160–169. https://doi.org/10.1016/j.cmet.2011.01.003.

[30] Liu JQ, Zhang L, Yao J, Yao S, Yuan T. AMPK alleviates endoplasmic reticulum stress by inducing the ER-chaperone ORP150 via FOXO1 to protect human bronchial cells from apoptosis. Biochemical & Biophysical Research Communications. 2018. 497:564–570. https://doi.org/10.1016/j.bbrc.2018.02.095.

[31] Woodworth-Hobbs ME, Hudson MB, Rahnert JA, Zheng B, Franch HA, Price SR. Docosahexaenoic acid prevents palmitate-induced activation of proteolytic systems in C2C12 myotubes. J Nutr Biochem. 2014;25:868–874. https://doi.org/10.1016/j.jnutbio.2014.03.017.

[32] Cacicedo JM, Benjachareowong, S, Chou E, Ruderman NB, Ido Y. Palmitate-induced apoptosis in cultured bovine retinal pericytes: roles of NAD(P)H oxidase, oxidant stress, and ceramide. Diabetes. 2005;54:1838–1845. https://doi.org/10.2337/diabetes.54.6.1838.

[33] Coll T, Jove M, Rodriguez-Calvo R, Eyre E, Palomer X, Sanchez RM, et al. Palmitate-mediated downregulation of peroxisome proliferator-activated receptor-gamma coactivator 1alpha in skeletal muscle cells involves MEK1/2 and nuclear factor-kappaB activation. Diabetes. 2006;55:2779–2787. https://doi.org/10.2337/db05-1494.

[34] Chen C, Kassan A, Castañeda D, Gabani M, Choi SK, Kassan M. Metformin prevents vascular damage in hypertension through the AMPK/ER stress pathway. Hypertens Res. 2019;42:960–969. https://doi.org/10.1038/s41440-019-0212-z.

[35] Varshney R, Varshney R, Mishra R, Roy P. Kaempferol alleviates palmitic acid-induced lipid stores, endoplasmic reticulum stress and pancreatic β-cell dysfunction through AMPK/mTOR-mediated lipophagy. J Nutr Biochem. 2018:212–227. https://doi.org/10.1016/j.jnutbio.2018.02.017.

[36] Yang M, Zhang D, Zhao Z, Sit J, Saint-Sume M, Shabandri O, et al. Hepatic E4BP4 induction promotes lipid accumulation by suppressing AMPK signaling in response to chemical or diet-induced ER stress. The FASEB Journal. 2020;34:13533–12547. https://doi.org/10.1096/fj.201903292RR.

[37] Zhang J, Wang Y, Bao C, Liu T, Huang J, Li J. Curcuminloaded PEGPDLLA nanoparticles for attenuating palmitateinduced oxidative stress and cardiomyocyte apoptosis through AMPK pathway. International Journal of Molecular Medicine. 2019;44:672–682. https://doi.org/10.3892/ijmm.2019.4228.

[38] Salvadó L, Barroso E, Gómez-Foix AM, Palomer X, Michalik L, Wahli W, Vázquez-Carrera M. PPARβ/δ prevents endoplasmic reticulum stress-associated inflammation and insulin resistance in skeletal muscle cells through an AMPK-dependent mechanism. Diabetologia. 2014;57:2126–2135. https://doi.org/10.1007/s00125-013-2867-3.

[39] Kwak HJ, Choi HE, Cheon HG. 5-LO inhibition ameliorates palmitic acid-induced ER stress, oxidative stress and insulin resistance via AMPK activation in murine myotubes. Scientific Reports. 2017;7:5025. https://doi.org/10.1038/s41598-017-05346-5.

[40] Yang L, Sha H, Davisson RL, Qi L. Phenformin activates the unfolded protein response in an AMP-activated protein kinase (AMPK)-dependent manner. Journal of Biological Chemistry. 2013;288:13631–13638. https://doi.org/10.1074/jbc.M113.462762.

